# Molecular profiling of lipid droplets inside HuH7 Cells with Raman micro-spectroscopy

**DOI:** 10.1101/2020.02.27.968701

**Authors:** Ashok Zachariah Samuel, Rimi Miyaoka, Masahiro Ando, Anne Gaebler, Christoph Thiele, Haruko Takeyama

## Abstract

Raman imaging has become an attractive technology in molecular biology because of its ability to detect multiple molecular components simultaneously without labeling. Two major limitations in accurately accounting for spectral features, viz. background removal and spectral unmixing, have been overcome by employing a modified and effective routine in multivariate curve resolution (MCR). With our improved strategy, we have spectrally isolated seven structurally specific biomolecules without any post-acquisition spectral treatments. Consequently, the isolated intensity profiles reflected concentrations of corresponding biomolecules with high statistical accuracy. Our study reveals the changes in the molecular composition of lipid droplets (LD) inside HuH7 cells and its relation to the physiological state of the cell. Further, we show that the accurate separation of spectral components permits analysis of *in vivo* structural modification of molecules after cellular uptake. A detailed discussion is presented to highlight the potential of Raman spectroscopy with MCR in semi-quantitative molecular profiling of living cells.

## Introduction

Living cells are complex entities with organelles formed by collectively existing molecules performing specific functions. One such seemingly simple but dynamic cell organelle is lipid droplet. Lipid droplets, which are made up of lipid-rich core surrounded by monolayer of phospholipids, are generally thought to be storage organelles. ^1,2^ There is a great amount of interest in molecular profiling of LDs inside cells as more and more evidences reveal its functional role^3^ beyond storage of molecules.^4,5^ LDs play significant, but yet not completely understood, role in diseases; for instance, fatty liver disease, diabetes,^6^ hepatitis C virus (HCV) infection,^7^ and cancer.^8^ LDs have been shown to play significant roles in viral assembly during virus replication,^9^ as storages sites for specific histones,^10^ mediators in transcription,^11^ phosphatidylcholine synthesis^12^ etc. These studies indicate a larger role LDs play in sustaining life. Molecular profiling of LDs is hence an extremely important aspect of molecular biology.^13^

Fluorescence microscopy is extensively used in molecular biology for investigating distributions of molecules and organelles, including LDs, inside single cells. Lipid-selective fluorescent molecules are generally used to light-up LDs inside cells (Nile Red ^14^ and BODIPY^15^). However, a study of distribution of several different molecules inside LDs would require each of them to be selectively labeled. In such efforts, it is necessary to avoid lager molecular structural alterations of the probe-molecule such that the biological functional equivalence of the labeled species is largely unaffected compared to its natural analogue. ‘Alkyne tagging’^16,17^ has been demonstrated as one of the smallest possible ‘tagging’ methods for subsequent fluorescent labeling. Supporting this argument, *in vivo* esterification of alkyne cholesterol in glioblastoma cells were found to proceed efficiently, equivalent to ‘*untagged’* cholesterol.^16^ Derivatives of PC lipids with strategically extended conjugation also serve as fluorescent molecules with minimal structural alteration for single cell imaging.^18^ However, a critical evaluation of biological behavior of labeled and native molecules inside a cell is very important in the light of extensive use molecular labeling methodology in molecular biology and medicine. This would require selective detection of these derivatives when they are simultaneously present inside the same cell. Earlier such attempts were often unsuccessful.^16^ We have fed cells with alkyne-tagged cholesterol both to evaluate accuracy of the analytical method and to test the biological equivalence of natural and tagged cholesterol derivatives.

Raman micro-spectroscopy is well suited for molecular profiling since it can detect multiple molecules simultaneously. Molecules such as proteins, lipids, cholesterol, nucleic acid etc. show distinguishable vibrational spectrum permitting molecular imaging of living cells.^19^ However, linear Raman imaging technique has often been criticized for the inferior signal strength compared to the nonlinear analogues such as coherent anti-stokes Raman spectroscopy (CARS) and stimulated Raman scattering (SRS).^20^ But it should be noted that, in order to get full spectral information, CARS and SRS employ high power pulsed lasers and are shown to be detrimental to the biological constituents.^21-23^ Use of a narrowband laser pulse instead would allow rapid image acquisition but at the cost of multiplex ability.^22^ Further, in CARS, the nonlinear polarization in an inhomogeneous system (cells) along with non-resonant background results in spectrum considerably different from the corresponding Raman spectrum.^22^ Local environments, number, shape and size of scatterers also affect CARS spectrum and intensity leading to difficulty in directly estimating the composition of different molecular species in biological cells.^23^ Therefore, nonlinear techniques have limitations when it comes to molecular concentration profiling of cells. However, remarkable rapid imaging capabilities of these techniques provide useful information on distribution of lipids in biological cells and tissues.^20, 25-27^

Ease of multiplexing is an advantage of Raman spectroscopy over fluorescence and nonlinear Raman techniques. At the same time, due to multiplexing, a composite Raman spectrum of several molecular components is generated from each focal spot during imaging. Therefore, multiplex Raman imaging require efficient data analysis techniques that effects separation of spectral components and their concentration profiles from the composite spectrum. Multivariate curve resolution by alternating least squares (MCR-ALS) combined with singular value decomposition (SVD)^28-31^ is an appropriate multivariate analytical technique that separates interpretable spectral components and their concentration profiles. The efficacy of the method has been demonstrated in earlier studies including several of our own.^28-31^ However, when applied to biological systems, this technique encounters a serious problem from highly fluctuating background. This often get manifested as signal-signal, signal-background mixing and limits spectral unmixing. Therefore, undesirable arbitrary baseline correction and other post-acquisition spectral modifications are routinely performed. In this study we demonstrate a modified MCR routine that allows better signal separation from broad backgrounds with highly improved spectral unmixing. Consequently, we could obtain intensity profiles reflecting statistically accurate compositions of biomolecules that reflected the biological features of the cells and organelles.

In the present investigation, we have applied spontaneous Raman spectroscopy and the improved MCR-ALS routine for molecular profiling of individual lipid droplets inside HuH7 human liver cells. LDs in each cell show small but considerable variations in the concentrations of molecular constituents but a drastic change was observed upon changing the cell-culturing condition. Through detailed analysis, we provide explanation for these changes based on statistically significant variations in the molecular composition of the cell. A detailed discussion of the data analysis to emphasize the potential of Raman-MCR technique in analyzing biological systems is provided in the manuscript.

## Results and Discussion

HuH7 human liver cells were cultured in laboratory using DMEM medium in a chamber with glass bottom (see Supplementary information S1 for experimental details). Two different feed conditions (table 1) were used in the present study: 1) OL-feed condition - culture medium containing oleate and alkyne cholesterol 2) CH-feed condition - culture medium containing cholesterol and alkyne cholesterol. Six different HuH7 cells were analyzed in the present study: three cells under each feed condition.

**Table 1.** Different feed conditions used for the study.

| Name | Second day |  | Third Day |  |  | Cells |
| --- | --- | --- | --- | --- | --- | --- |
| | Cholesterol*<br>(μM) | Oleic<br>Acid<br>(μM) | Cholesterol<br>(μM) | Oleic<br>Acid<br>(μM) | Alkyne Cholesterol*<br>(μM) | |
| **OL-feed** |  | 50 |  | 50 | 50 | 1 to 3 |
| **CH-feed** | 50 |  | 50 |  | 50 | 4 to 6 |

\* The oleic Acid, cholesterol and alkyne cholesterol in the culture medium is in the free form and they get enzymatically esterified after cellular uptake (see the discussion).

A white light image of one of the HuH7 cells containing LDs is provided in figure 1a. LDs are discernable in the figure because of their globular appearance (slight distortion in shape may be due to cell fixing and placing of a cover slide). The triple bond stretching vibration of alkyne cholesterol appears in an isolated region (2119 cm^-1^) of the Raman spectrum allowing an accurate estimation of its distribution in the cell. The Raman image thus obtained (Figure 1c) has high resemblance to the LDs found inside the cells (Figure 1a). This indicates that alkyne cholesterol is highly concentrated in LDs. We found that the perimeter obtained after ∼20% intensity thresholding agrees well with the LD boundaries seen in the white light image (Figure 1b). Thus, we could estimate the total area occupied by LDs in each cell. Our analyses indicate a higher proportion (total area) of LD in cells cultured under OL-feed condition (∼2 times) compared to CH-feed.

**Figure 1.**
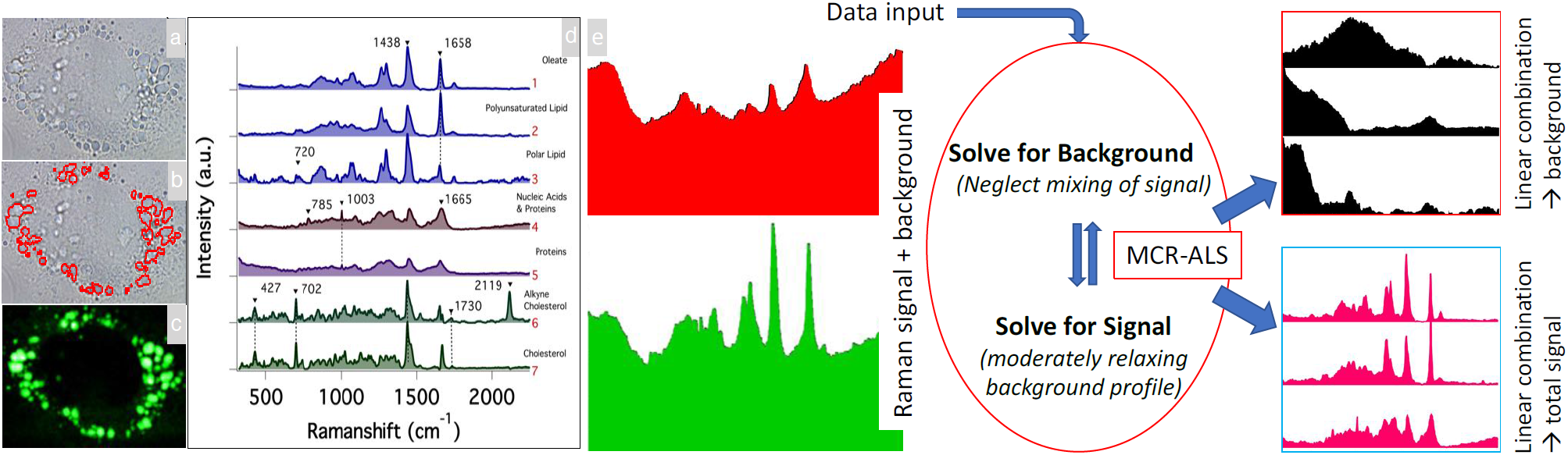
a) White light Image of the cell. Lipid droplets can be identified as globules. b) LD boundaries. c) Spatial distribution of intensity of alkyne bond stretching mode (2119 cm^-1^) of alkyne cholesterol. d) Significant spectral components separated in the MCR-ALS analysis. We have assigned them as, 1 - Oleate, 2 – Polyunsaturated lipid, 3 – polar lipid, 4 - Nucleic Acid and protein, 5 – Protein, 6 - Alkyne cholesterol, 7 - Cholesterol. There were also broad background spectral components without any significant biological spectral features. Full list of spectral components is given in supplementary information S2. The accuracy of the analysis is clear from the negligible residual (supplementary information S3). e) A pictorial depiction of modified MCR-ALS routine used in this study. A linear combination of the separated profiles (background and signal) gives total Raman spectrum.

Presence of an isolated C≡C Raman band allowed direct imaging of alkyne cholesterol distribution. However, the fingerprint region <2000 cm^-1^ of the Raman spectrum is highly overlapped. Hence, quantitatively extracting band intensities of other biological components is not easy. Multivariate curve resolution (MCR) is a useful method to effect spectral unmixing.^28-30^ Since the weak Raman signals are often mixed with highly fluctuating strong background signals, arriving at proper MCR solution is not easy. We have addressed this issue by avoiding all post-acquisition spectral modification and solving for backgrounds or Raman signals focusing on one at a time (codes were written in *Python)*. In the new routine, starting with SVD guess for number of components, we keep modifying the number of components (**k**) in MCR (Supplementary information S1) until broad signals separate out. However, in doing so Raman spectral components often gets scrambled, which we neglect in the initial cycles. Then, broad spectral components identifiable as background were fed into a new MCR cycle and solved for Raman signals. In this cycle, a) **k** value is changed until a reasonable solution is obtained with constrained background signal components and then b) the constraints on the background signals are moderately relaxed and solutions are further optimized. These steps are repeated until a stable solution is obtained such that thereafter no appreciable modifications of the spectral profiles occur. At this stage Raman images are inspected to confirm acceptability. Known background signals (e.g. from glass) are given as initial inputs to speed up the process of background separation. This modified MCR-ALS routine was applied (cartoon depiction Figure 1) to a large dataset of 91,220 Raman spectra from six different HuH7 cells cultured under two different feed conditions. This big-data was used as a single matrix (A; see supplementary information S1 - MCR-ALS section) to perform MCR-ALS based dimensionality reduction to retrieve relevant spectral components and the corresponding concentration matrix. The accuracy of the presented model is evident from the negligible MCR-ALS residual (Supplementary information S3). The result of MCR-ALS is provided in figure 1d. Seven significant molecular constituents were separated in the analysis.

### Analysis of lipids

Details of assignment of spectral components is given in supplementary information S2. MCR components 1, 2 and 3 are lipid spectra (Figure 1d) and we have assigned them as follows. Component-1 as oleate, component-2 as polyunsaturated lipid and component-3 as polar lipid. In the lipid spectra the intensity of the Raman band at 1658 cm^-1^ indicates extent of unsaturation (C=C) in the lipid.^32^ Polyunsaturated lipid has significantly higher intensity at 1658 cm^-1^. Polar lipid component has relatively lesser degree of unsaturation and a band at around 720 cm^-1^ (choline group) indicating it polar nature (e.g. phosphatidylcholine).^32^

MCR-ALS results also gives spatial distribution of individual spectral components (H matrix; see supplementary information S1). Spatial distribution (Raman images) of the three lipid-species are provided in figure 2. Oleate and polyunsaturated lipid can be found localized in LDs. Polar lipids are found distributed in cytosol medium. In order to better understand the spatial correlation between individual lipid distributions, pixel-to-pixel intensity correlation was performed (images display contrast). Pearson correlation plots^33^ for lipids are shown in the figure 2 (*see* also supplementary information S4). Data points occupying diagonals indicate a perfect correlation in Pearson’s intensity-intensity correlation plots.^33^ The corresponding Pearson’s correlation (PC) coefficients are given as inset. Perfectly oppositely correlated images give a PC value of −1 while PC=1 indicates a perfect positive correlation.^33^ It can be seen that distributions of oleate and polyunsaturated lipid are spatially well correlated (Figure 2). Negative correlation between oleate and polar lipids (Figure 2), on the other hand, reconfirms the spectral assignment. The difference in polarity leads to their mutually exclusive spatial distribution. Further, a good correlation between distributions of alkyne cholesterol and oleate is also found (supplementary information S5; Table S1). Our data indicates that cores of LDs are rich in non-polar lipids and sterol esters.

**Figure 2.**
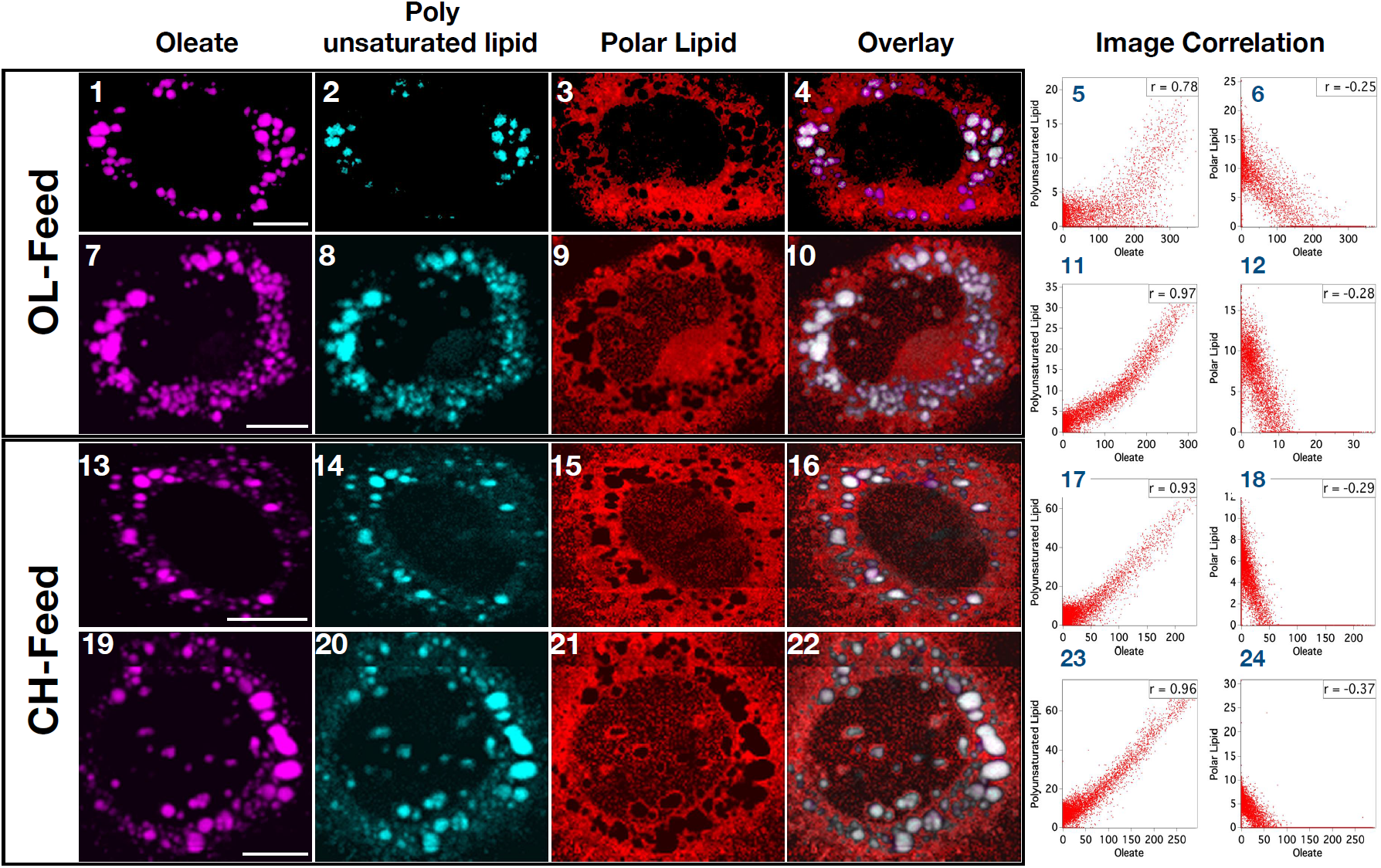
Distribution of lipids in HuH7 cells. Overlay of lipid distributions are provided in the fourth column. Oleate and polyunsaturated lipid are almost entirely localized in LDs while polar lipid is distributed in cytosol medium. Images from 4 representative cells are shown here (cells 1, 2, 4 and 5; *see* also supplementary information S6). Pearson image correlation plots for lipids (*see* also supplementary information S4) are shown in the last two columns. Correlation coefficients are indicated as insets in each plot. Scale bar 10 μm.

### Analysis of proteins and nucleic acids

MCR components 6 and 7 are nucleic acids and protein respectively (supplementary information S2).^34^ MCR component 6, has a spectral marker band for nucleic acids at 785 cm^-1^ (phosphodiester vibrational mode)^35^ in addition to prominent protein bands. It means there are some protein species that coexist with nucleic acids. The spatial distribution of these two components in different cells are shown in figure 3. Proteins are distributed in cytosol (Cyan) while nucleic acids (Yellow) are rich in the nuclear region. Ring like and patchy appearance of proteins are also noticed on LD surfaces (figure 3) indicating proteins associated with LDs.^1, 13^ The technique, however, doesn’t have adequate spatial resolution to investigate proteins exclusively attached to the LD monolayer surface. ^36^ Hence, we think that the estimate of Raman intensities of proteins at LD gives an overestimate of its concentration (Spatial resolution is ∼300 times larger than the LD surface membrane dimension). But the relative variation in the observed protein intensity may reflect the trend of LD-to-LD composition variation. However, a possibility of aggregation (not membrane bound) of protein around LDs may not be completely ruled out (Figure 3-1). Further, dense aggregates of proteins and nucleic acids are observed within the nuclear region (Figure 3; Bright yellow). Based on the size of these aggregates and earlier reports^37^, we assign these as nucleoli. Detailed study is required to completely understand the nature of these aggregates.

**Figure 3.**
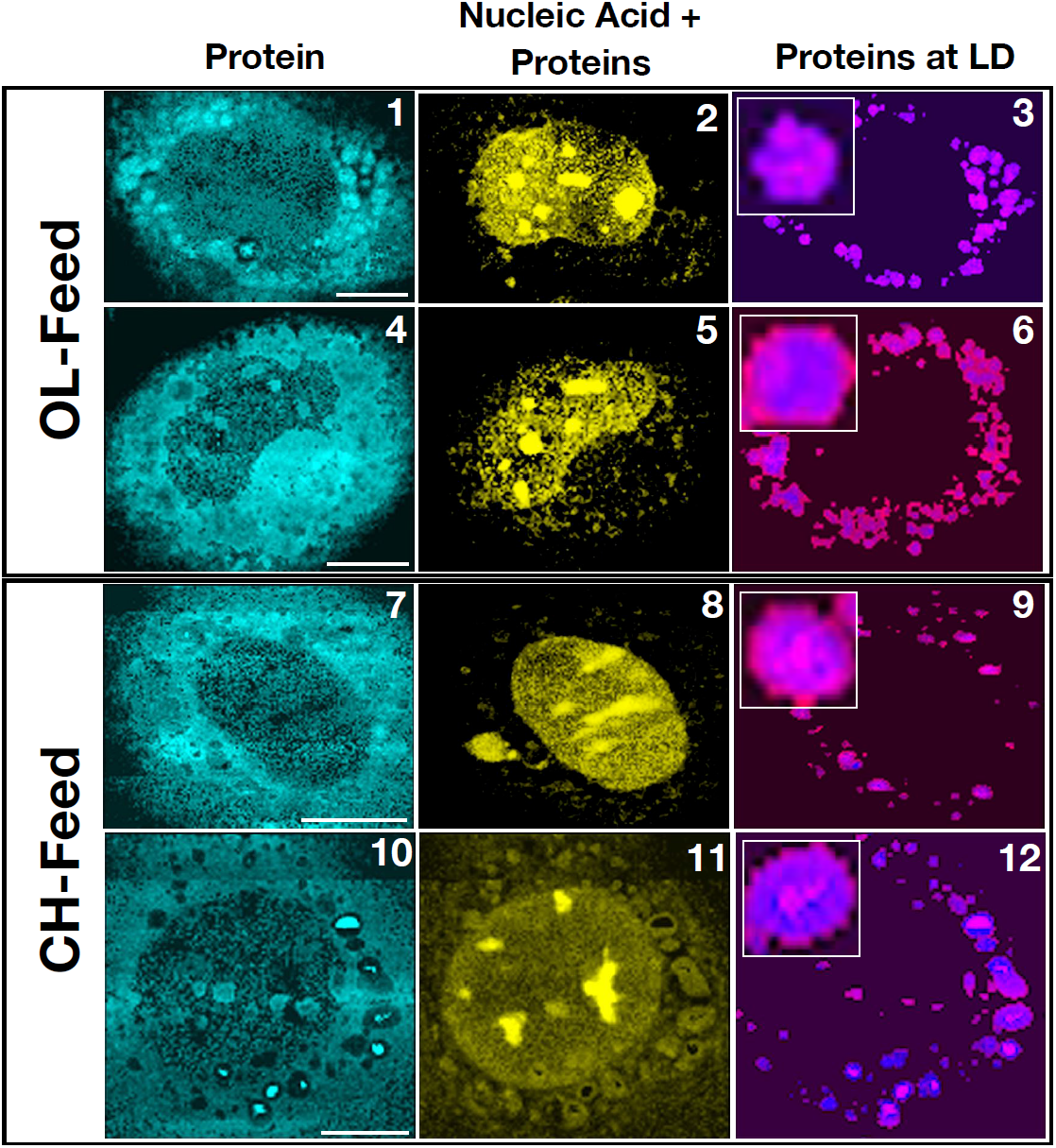
Distribution of Proteins (Cyan) and a mix of nuclear proteins + nucleic acids in HuH7 cells. Proteins colocalized with LD are shown in the fourth column. Location of LDs (blue) were determined as described in the text. Images from 4 representative cells are shown here (see also supplementary information S6). Scale bar 10 μm.

### Analysis of sterols

Raman spectra of cholesterol and alkyne cholesterol got separated (Figure 1d) in the MCR analysis. This permits a comparison of their relative spatial distribution simultaneously. A band at 1730 cm^-1^ in these spectral components indicates that the sterols are present as esters in the cells.^32^ That is, both the sterol derivatives from the feed get enzymatically converted into the corresponding esters *in vivo*. Hereafter the names cholesterol and alkyne cholesterol indicate corresponding ester forms unless otherwise specified. When referring to the non-esterified sterols “free form” will be provided in parenthesis after corresponding sterol names.

As discussed earlier, alkyne tagging is one of the simplest structural modifications to a biological molecule such that it behaves identical to the natural analogue. This tag can then be used to attach a fluorophore to visualize molecules using fluorescence spectroscopy.^16-18^ In the present study we intended a critical evaluation of biological behavior of cholesterol and alkyne cholesterol.

Several possible scenarios can be expected to occur during the cell culture (Table 1): a) cells uptake the sterols (free form) irrespective of the structural modification, but alkyne cholesterol is kept isolated in LDs, b) when both cholesterol (free form) and alkyne cholesterol (free form) are simultaneously provided in the feed, a preferential uptake of cholesterol (free form) by the cells occur, or c) the cells treats both cholesterol and alkyne cholesterol equally. In order to successfully evaluate which of these actually occurs, it is important to selectively detect these structural analogues in the same cell under the same conditions. Under OL-feed condition, only alkyne cholesterol (free form) is provided in the culture medium and under this feed condition any cholesterol detected inside the cell is the cholesterol naturally existing in the cells. Analysis of Raman images of the cells cultured under this condition will then permit evaluating whether alkyne cholesterol is completely sequestered in LDs or not. Under the CH-feed condition, on the other hand, both the sterols (free form) are provided in the culture medium such that their uptake by the cell and spatial distribution inside the cell can be compared. Since MCR-ALS analysis has separated the spectra of these sterol derivatives, we could perform such a comparative study accurately.

The spatial distributions of alkyne cholesterol (green) and cholesterol (blue) are given in figure 4. At first glimpse it appears that sterol derivatives have different spatial distributions (figure 4-1 & 2). Under OL-feed condition, cholesterol is also found outside LDs while alkyne cholesterol is found rich inside LDs (figure 4-1 & 2). On the other hand, in the Raman images of the cells cultured under CH-feed, both the sterols can be found co-localized in LDs (figure 4-11 & 12). It should be noted, however, that these images represent contrast of Raman intensities. That is, high intensity of sterol esters inside LDs makes their distribution outside LDs less apparent in the images. That is depending on the concentration of sterols in LDs, the images may appear different and may not completely depict the true scenario. In order to gain a better understanding of their relative spatial correlation, sterol distributions inside and outside LDs must be separately analyzed. Such an analysis was performed and the distributions of sterols outside LDs are given in Figure 4 (last two columns). Remarkable similarity between the distributions can be seen from the plots. Distribution of sterol derivatives outside LD shows high correlation with an average Pearson Correlation coefficient of 0.79 (Table S1). Thus, we show that spatial distribution of these sterols inside HuH7 cells are identical within the experimental limitation of spatial resolution. Further, our results indicate that the distribution of cholesterol inside LDs depends on the feed condition. Absence of cholesterol (free form) in the OL-feed leads to its poorer concentration inside LDs. However, both sterols are abundantly present inside LDs (Pearson Correlation coefficient >0.9; Table S1) in cells cultured under CH-feed because both are provided in the feed.

**Figure 4.**
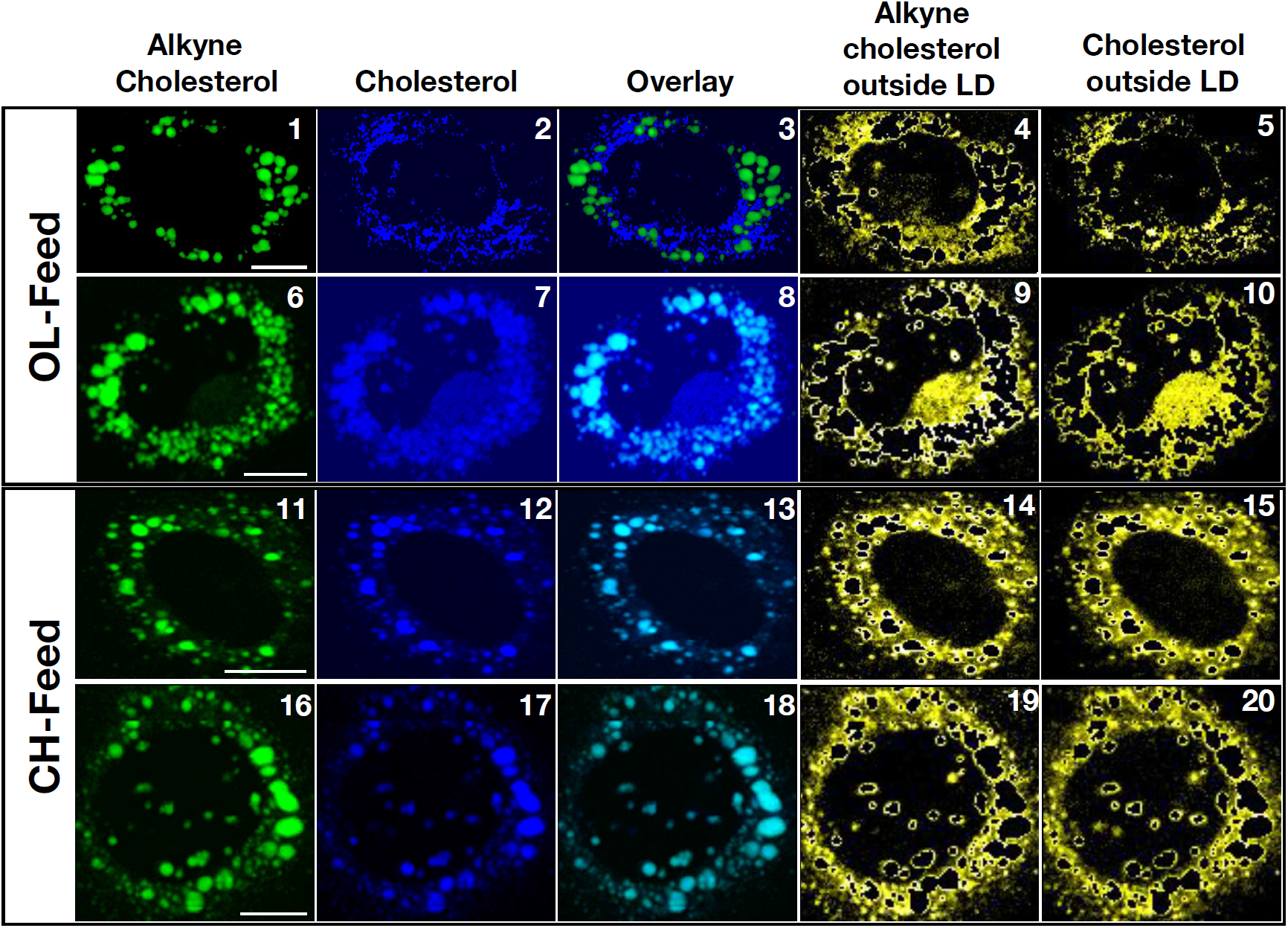
a) Distribution of cholesterol and alkyne cholesterol in HuH7 cells. The distribution of cholesterol outside LDs (yellow) are shown in the last two columns for cholesterol and alkyne cholesterol respectively. Identical distribution can be noted. Images from 4 representative cells are shown here (see also supplementary information S6). Scale bar 10 μm.

### Total sterol uptake by HuH7 cells

*In vivo* esterification and identical spatial distribution inside the cells point to the most-probable biological equivalence of these sterol derivatives. However, this should be further verified by comparing their total uptake by the cells. Total Raman intensities of MCR spectral components (ΣH; supplementary information S1) reflects the concentration of these components in cells. Average values of percent composition of sterol components (e.g. %Alkyne cholesterol; *see* Supplementary information S1) inside the cells are shown in figure 5A. Under OL-feed condition, ∼90% of the total sterol in LDs is alkyne cholesterol. This is because, under this culture condition, only alkyne cholesterol (free form) is provided in the culture medium (Table 1). Cholesterol observed under this condition is the ones naturally existing in the cells. On the other hand, under CH-feed condition 2:1 cholesterol to alkyne cholesterol ratio is observed (Figure 5A). It agrees with the composition of cholesterol (free form) and alkyne cholesterol (free form) in the culture medium (2:1; see Table 1). The total Raman intensities of sterols per cell (Figure 5B) is nearly the same (p>0.05; no statistical difference). We have shown that, 1) both sterol derivatives (free form) taken up from the medium gets enzymatically esterified inside HuH7 cells, 2) the spatial distribution of these sterols are identical and 3) the total uptake by the cell is in accordance with the feed composition. Thus, we infer that attaching a triple bond at the end of alkyl-chain of cholesterol doesn’t considerably alter its cellular behavior. This also confirms that the Raman intensities observed in our experiments can account for the molecular composition of the cell (also LDs).

**Figure 5.**
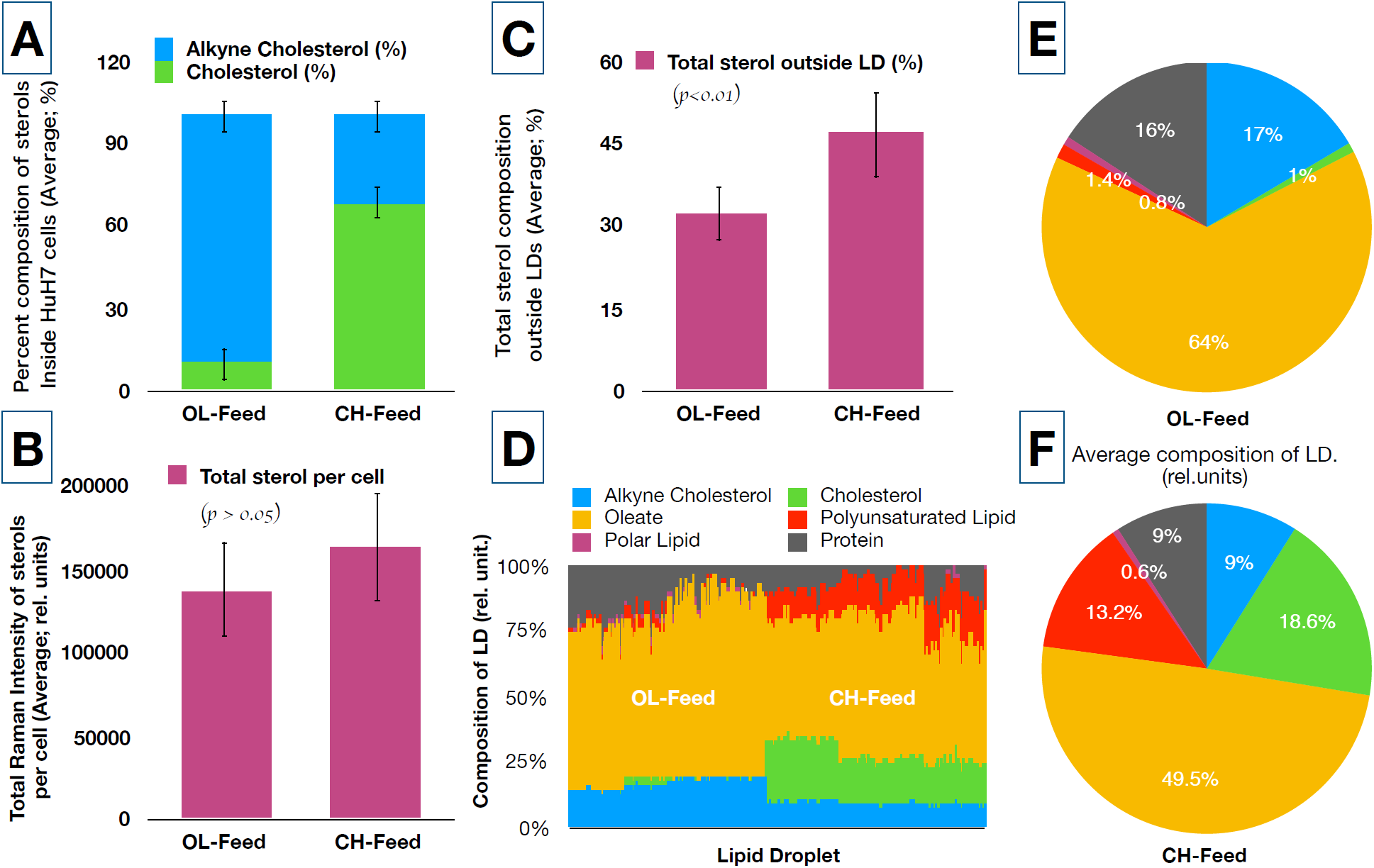
A) Percent compositions (Raman intensities) of alkyne cholesterol and cholesterol inside HuH7 cells (relative to total sterol) under two different feed conditions. The error bars represent standard deviations. B) Total Raman intensities (sterol composition) inside the whole cell. Statistically insignificant differences from the student t-test (p>0.05); total sterol uptake is comparable under two feed conditions. C) Total sterol composition outside LDs under two feed conditions. Statistically significant (p<0.01) excess composition of sterols found outside LDs under CH-feed. D) Molecular composition of individual LDs under two culture conditions. The height of each colored bar represents area normalized Raman intensity inside each LD (in % scale). The pie charts E and F shows average value of area normalized total Raman intensities of individual molecular components under two different feed conditions. The Raman intensities are proportional to concentration and can be converted to absolute concentrations by knowing Raman cross-sections.

### Molecular Profiling of LDs in HuH7 cells

Quantifiability of the MCR concentration profiles evident from the analysis discussed above gives confidence to analyze molecular concentration profiles inside each LD present in HuH7 cells. To do this we have estimated the total intensity of each molecular component inside each LDs (229 individual LDs) from the corresponding concentration matrices (H). Raman intensities within the LD areas are only used in this analysis. Area normalized compositions of LDs are shown in figure 5D, where each vertical bar plot represents composition of one LD. Average value of these compositions under OL and CH feed composition are given in figure 5E & F respectively. Under OL-feed condition an LD has on an average 17% alkyne cholesterol and 1% cholesterol (94:6). Similarly, under CH-feed condition average sterol composition of LD changes to 9% alkyne cholesterol and 19% cholesterol in agreement with the 2:1 feed composition. About 64% of the total composition of LDs is lipid (under both feed conditions). Protein composition is found to be different (16% and 9%) under two feed conditions. Since we believe these numbers are over estimated due to limited spatial resolution, it is not possible to interpret the observed differences. However, a considerable protein composition variation can be expected on LDs. Oleate composition reduces from 64% under OL-feed to 50% under CH-feed (Table 1). Importantly, the polyunsaturated lipid composition is significantly affected by the feed condition. Under OL-feed condition, only 1% of polyunsaturated lipid is observed inside LDs (Figure 5E). Under CH-feed condition, however, an order of magnitude increase in polyunsaturated lipid composition is observed (Figure 5F) inside each cell (Figure 5D). We find this behavior is strongly correlated to the total sterols found outside LDs (total sterol composition per cell is statistically similar; figure 5C): the total composition of sterols outside LDs (Figure 5C) increases significantly (p<0.01) under CH-feed (46%) compared to OL-feed (32%). This higher concentration, we believe, in turn triggers an inflammatory response (production of polyunsaturated lipids^38,39^) in the cell.^38-40^ Nearly half of the polyunsaturated lipid produced in the cell is found stored in LDs (Supplementary information S7). Thus, it is clear that the composition of LD changes depending on the physiological condition of the cell.

## Conclusions

We have demonstrated the effective use of Raman spectroscopy and MCR-ALS for the simultaneous imaging of multiple molecular species inside HuH7 cells. We have analyzed a big-data collection of 91,220 spectra from 6 different HuH7 cells. Our MCR model successfully explains all the spectral variations in the dataset (negligible residuals). Raman imaging could locate and quantify the proportion of LDs in HuH7 cells under different feed conditions; such a quantification has relevance in the disease conditions.^41,42^ Previous Raman imaging studies have also shown the capability of the technique to detect LDs in cells by mapping lipids. ^43-47^ However, molecular mapping of many different constituents of LD has never been demonstrated. In our study, seven different molecular species were identified. Three structurally different lipid species have been spectrally isolated, and the relevance of the corresponding relative compositions has been discussed. By simultaneously providing cholesterol (free form) and alkyne cholesterol (free form) in the culture medium, we could compare the biological behavior of the native and structurally modified sterol derivatives. We confirmed enzymatic esterification of sterols inside cells, identical spatial distribution and nondiscriminatory cellular uptake. Our data shows a high correlation between the concentration of sterols located outside-LDs in HuH7 cells and the over-expression of polyunsaturated lipids. We could clearly identify the changes caused by differences in the composition of culture medium and those caused by biological responses. Raman imaging with MCR-ALS has enormous potential to be used for molecular profiling.

## Acknowledgements

This work was supported by Grant-in-Aid for Scientific Research(S) (no. 17H06158).

## Author information

### Author notes

These authors contributed equally: Ashok Zachariah Samuel and Rimi Miyaoka.

### Contributions

A. Z. S conducted data analysis, image analysis, manuscript writing. R. M. conducted Raman imaging experiments. M. A. conducted MCR-ALS. A. Z. S. and R. M. contributed equally to the work. A. G. and C. T. provided the HuH7 cells for the study. H. T. and M. A. contributed to manuscript writing and discussion.

## Supplementary information

Supplementary information is available. SI includes details of experimental methods, full list of MCR-ALS spectral components, all Raman images of cells, MCR residuals plot, Pearson Plots and a table of correlation coefficients.

