## supplementary information for "Molecular profiling of lipid droplets inside HuH7 Cells with Raman micro-spectroscopy"

**Supplementary information S1**

**Experimental Section**

**Cell culture and Biological sample preparation**

**Collagen coating of the glass bottom chamber for cell culture:** Cell matrix type IV (Nitta Gelatin Inc., Osaka, Japan) containing collagen was diluted with 10% hydrochloric acid. 1mL of this diluted collagen solution was added into the glass bottom chamber so that the solution uniformly covers the surface. After a small waiting period, the solution was carefully removed. The chamber was then incubated for about 30-60 min at room temperature. Then the chamber was washed with the medium.

**Cell Culture:** HuH-7 (JCRB0403) cells were first cultivated in a culture flask using DMEM (Gibco, 11965092) medium at 37 °C. The culture was supplemented with 10% (v/v) lipid-free fetal bovine serum and 5 % (v/v) $CO_2$. The cells were then ($1\text{-}2\times10^5$ cells/mL) inoculated in a glass bottom chamber (Thermo) coated with collagen. After a day selected cells were subjected to two different feed conditions of OL-feed and CH-feed.

**Feed conditions:** In one feeding condition, named as **OL-feed**, 50μM oleic acid was fed on the second day. A mixture of 50μM oleic acid and 50μM alkyne cholesterol was fed on the third day. In another feed condition, named **CH-feed,** 50 μM cholesterol was fed on the second day instead of oleic acid. A mixture of 50 μM cholesterol and 50 μM alkyne cholesterol was fed on the third day.

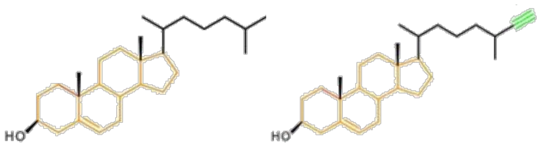


Cholesterol Alkyne Cholesterol

**Cell fixing:** On the fourth day, the cells were washed with PBS and 1mL PFA (4%) was added. It was then incubated for 30 min. PFA was then removed by washing with PBS followed by deionized water.

**Raman Spectroscopy**

Raman microspectroscopic imaging measurements were carried out with a laboratory-built confocal Raman microspectrometer. A 632.8 nm line of a He-Ne laser (HRP350-EC - HeNe Laser, THORLAB, USA) was used as the laser source. The laser beam was focused into the sample with a 100X (1.4 NA) objective lens (Plan Apo VC; Nikon Corporation, Tokyo, Japan) mounted on an inverted microscope (ECLIPSE Ti; Nikon Corporation, Tokyo, Japan). The back-scattered Raman light was collected with the same objective lens and measured with a spectrometer (MS3504i, 600 lines/mm; SOL Instruments, Ltd., Minsk, Republic of Belarus) and a CCD detector (Newton DU920-M; Andor Technology Plc., Antrim, UK). The laser power at the sample was 15 mW. A piezoelectric stage (custom-made; Physik Instrumente GmbH & Co. KG, Karlsruhe, Germany) was used to carry out Raman mapping measurements (0.3 µm step size). The exposure times were 0.2 to 0.3 s per point of the cell. Using indene as external calibration, Raman images are normalized (Normalized for laser intensity fluctuation between imaging experiments). Spectral, Raman image processing (interpolation, thresholding, area calculation etc.) and Pearson's correlation analyses were performed with IgorPro software. Images are plotted with Image J.

**MCR-ALS and SVD**

Multivariate curve resolution (MCR) by alternating least squares (ALS) is effective in identifying unique spectral components whose linear combinations constitutes the original data matrix. An original data matrix (A) can be decomposed into spectral components (W) and their concentration profiles (H) as given below.

$$A_{m,n} = W_{m,k}H_{k,n} + E_{m,n} \qquad \text{..........(S1)}$$

E is the error (or residual). In the present case, **A** (m × n matrix) consists of **n** spectra from different spatial locations each with **m** data points. In this method, the matrix **A** is decomposed to a matrix **W** with **m** × **k** dimensions, whose columns represent pure component spectra, and another matrix **H** with **k** × **n** dimensions, whose rows represent the intensity profiles of individual spectral components. MCR-ALS is performed iteratively by minimizing the error **E** such that the Frobenius norm $||A - WH||^2$ is minimized. Non-negativity constraints $W \geq 0$ and $H \geq 0$ are applied during the minimization procedure to obtain physically meaningful solutions. These constraints arise from the fact that neither the Raman spectra nor the concentration profiles will have negative values. In order to solve the MCR equation appropriately, the value of 'k' needs to be known or guessed. The number of independent spectral components can be obtained (at least as an initial guess value) by employing singular value decomposition (SVD). In SVD, original $\mathbf{m} \times \mathbf{n}$ matrix is decomposed into $U\Sigma V^T$ (U is m × m, Σ is m × n, and $V^T$ is n × n), where Σ represents the singular values (diagonal matrix). The number of dominant singular values gives the number of spectral components to be used for the MCR analysis. Remaining components necessarily will be noise. Therefore, SVD can also be used for noise reduction in the data with poor S/N ratio. This can be done by reconstructing the original data (e.g. **A**) as $U\Sigma V^T$ where singular values for all the noise components are kept zero.

The initial values of W and H matrices are also unknown to begin the calculations. We need to provide suitable initial guess values to optimize the decomposition. The SVD spectral components (U) or random numbers can be used as the initial guess spectra (initial W values). Then, the iterations to solve for W and H are performed alternately to arrive at the acceptable final solutions where residual is close to zero. It is possible that such a solution is not completely acceptable due to mixing of spectral components or incomplete removal of background etc. Under such situations, sparser solutions can be sought introducing regularization schemes such as L1 norm (Lasso regression; Tibshirani, 1996) or L2 norm (Ridge regression). L1 norm can be applied to H matrix or W matrix depending on situation as given below. If the concentration profiles show mixing of components L1(H) can be applied where as L1(W) regularization can be applied when mixing of spectral components are observed.

$$(W^T W + \alpha^2 E)H = W^T A \quad \text{..........(S2)}$$

$$(HH^T + \alpha^2 E)W = HA^T \quad \text{..........(S3)}$$

L2 norm can similarly be applied to H matrix or W matrix depending on situation as follows.

$$(W^T W + \beta^2 I)H = W^T A \quad \text{..........(S4)}$$

$$(HH^T + \beta^2 I)W = HA^T \quad \text{..........(S5)}$$

The optimized H matrix can be plotted as a 2D image representing the spatial distribution and the spectrum can be used to identify the chemical component.

**Calculating values in figure 5**

**Figure 5A**

Percent composition of cholesterol inside HuH7 cells =

$$\frac{\mathbf{100}\ \sum_{cell}\ H_{Cholesterol}}{\left(\sum_{cell}\ H_{Cholesterol} + \sum_{cell}\ H_{Alkyne\ Cholesterol}\right)} \quad \text{..........(S6)}$$

Percent composition of alkyne cholesterol inside HuH7 cells =

$$\frac{\mathbf{100}\ \sum_{cell}\ H_{Alkyne\ Cholesterol}}{\left(\sum_{cell}\ H_{Cholesterol} + \sum_{cell}\ H_{Alkyne\ Cholesterol}\right)} \quad \text{..........(S7)}$$

**Figure 5B**

Total Raman intensity of sterols per cell =

$$\left(\sum_{cell}\ H_{Cholesterol} + \sum_{cell}\ H_{Alkyne\ Cholesterol}\right) \quad \text{..........(S8)}$$

**Figure 5C**

Total sterol Composition outside LDs=

$$\frac{\mathbf{100}\left\{\left(\sum_{cell}\ H_{Cholesterol} + \sum_{cell}\ H_{Alkyne\ Cholesterol}\right) - \left(\sum_{LD}\ H_{Cholesterol} + \sum_{LD}\ H_{Alkyne\ Cholesterol}\right)\right\}}{\left(\sum_{cell}\ H_{Cholesterol} + \sum_{cell}\ H_{Alkyne\ Cholesterol}\right)}$$

..........(S9)

**Figure 5D**

Each vertical bar represents composition of one LD in an HuH7 cell. Each color code is the percent composition of one spectral component. For instance, the composition of oleate is calculated as follows. Since nucleic acids could not be separated as a pure component in the analysis, it was not included in the LD composition analysis. However, we found that the estimated compositions are not significantly affected by it.

Composition of oleate in one LD of area $A_{LD}$ =

$$\frac{\mathbf{100}\left(\sum_{LD}\ \boldsymbol{H}_{Oleate}\right)}{\boldsymbol{A}_{LD}\left(\sum_{LD}\left(\boldsymbol{H}_{Oleate}+\boldsymbol{H}_{Polyunsaturated\ Lipid}+\boldsymbol{H}_{Cholesterol}+\boldsymbol{H}_{Alkyne\ Cholesterol}+\boldsymbol{H}_{Polar\ Lipid}+\boldsymbol{H}_{Protein}\right)\right)}$$

………..(S10)

Compositions of other components were similarly calculated. More than 100 LDs were analyzed from cells cultured under each feed conditions. An average value of composition is plotted as a pie chart and is shown in figure 5E & F.

Summation represents adding Raman intensities at relevant pixels inside LD or cell as indicated. Reported numbers are average values.

**Supplementary information S2**

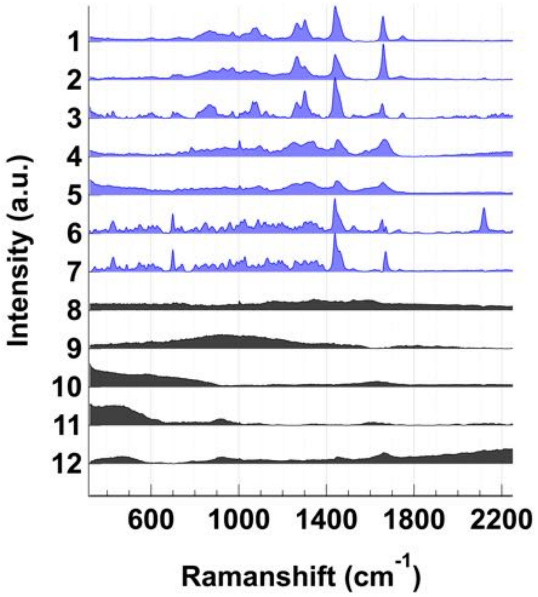


Figure S1. Complete list of MCR spectral components and background components (gray).

**Spectral Assignments**

In MCR components 1, 2 and 3 (Figure S1) a strong band at 1440 cm$^{-1}$ corresponding to -CH2-scissoring mode appears along with prominent bands at 1305 cm$^{-1}$ (CH- bending) and 1080 cm$^{-1}$ (C-C stretching rich in gauche conformation along the carbon chain) are observed. These are characteristic vibrations of alkyl chains and hence these components are lipids. Further, band at 1745 cm$^{-1}$ indicates ester carbonyl group of the corresponding ester group. The band at 1658 cm$^{-1}$ is due to unsaturation (C=C) in the alkyl chain. The intensity of this band indicates extent of unsaturation in the lipid. Component_2 has a higher intensity at 1658 cm$^{-1}$ compared to component _1 and component _3 indicating largest degree of unsaturation among the three lipids. Therefore, we assign component_2 as polyunsaturated lipid. A close similarity of component _1 spectral profile to oleate was confirmed by comparing with oleate Raman spectrum (*see* also Czamara, K., et al. J. Raman Spectrosc. 2015, 46, 4). Component _3 spectrum has relatively lower degree of unsaturation and a band at around 720 cm$^{-1}$, which indicates presence of choline functional group. Therefore, we assign component _3 to polar lipids (e.g. phosphatidylcholine).

MCR component 5 shows prominent protein spectral features such as, strong amide-I (peptide backbone vibration) at 1665 $cm^{-1}$, C-H deformation mode at 1450 $cm^{-1}$, tryptophan Cα-H deformation at 1340 $cm^{-1}$, phenylalanine at 1003 $cm^{-1}$. Therefore, MCR spectral component 7 represents proteins in the cell. MCR component 4, on the other hand, has an additional spectral marker band for nucleic acids at 785 $cm^{-1}$ (phosphodiester vibrational mode in addition to common protein spectral features) and hence it has been assigned to a mixture of nucleic acids and proteins.

Component_7 and component_8 shows characteristic vibrations of sterol ring below 1000 $cm^{-1}$, with prominent bands at 427 $cm^{-1}$ and 702 $cm^{-1}$. Further a strong band at 1438 $cm^{-1}$ corresponding to $CH_2$- bending mode and 1670 $cm^{-1}$ marker band corresponding to C=C stretch can be observed. Therefore, we assign these two components as sterols. Further, a band at 2119 $cm^{-1}$ (CC triple bond stretch) distinguishes component_7 from 8. Thus component_7 has been alkyne assigned as alkyne cholesterol and component 8 as cholesterol. Further, a small band at 1730 $cm^{-1}$ (ester C=O stretching) in these MCR components indicates that the sterol derivatives are present as esters.

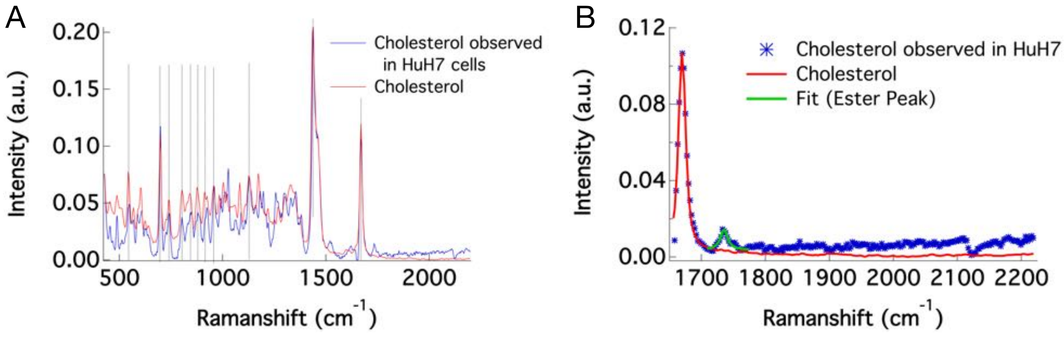


Figure S2. A) comparison of Raman spectra of standard cholesterol and cholesterol observed in cells. B) zoomed plot.

**Supplementary information S3**

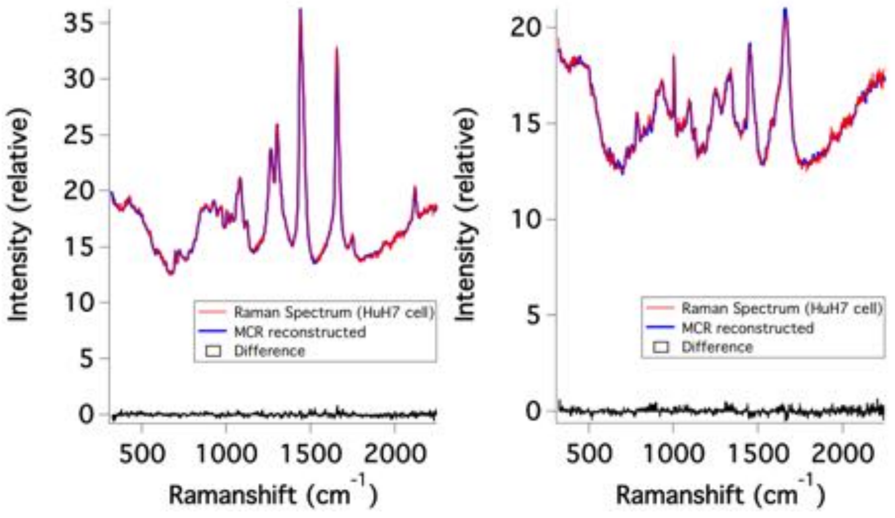


Figure S3. Residuals in the MCR-ALS analysis.

**Supplementary information S4**

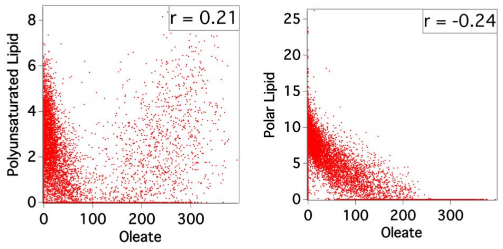


**Figure S4(a).** Pearsons's correlation between lipid images for a cell (cell-3; Cultured under OL-feed condition) which was not given in the main text.

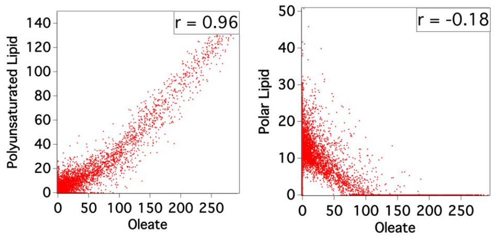


**Figure S4(b).** Pearson's correlation between lipid images for a cell (Cultured under CH-feed condition), which is not given in the main text.

**Supplementary information S5**

Table S1. Pearson's correlates coefficients (PC) for different pairs of Raman images.

| **Correlation Pair** | **Pearson's Correlation Coefficient for cells** | | | | | |
|---|---|---|---|---|---|---|
| | **1** | **2** | **3** | **4** | **5** | **6** |
| **Lipid 1 & 2** | 0.78 | 0.97 | 0.21 | 0.93 | 0.96 | 0.96 |
| **Lipid 2 & 3** | -0.12 | -0.28 | 0.42 | -0.17 | -0.18 | -0.13 |
| **Lipid 1 & 3** | -0.25 | -0.35 | -0.24 | -0.29 | -0.37 | -0.18 |
| **Alkyne Cholesterol & lipid 1** | 0.99 | 0.99 | 0.99 | 0.99 | 0.99 | 0.98 |
| **Cholesterol & alkyne cholesterol (Outside LD)** | 0.63 | 0.79 | 0.61 | 0.96 | 0.92 | 0.83 |
| **Cholesterol & alkyne cholesterol (Inside LD)** | 0.11 | 0.95 | 0.49 | 0.99 | 0.99 | 0.99 |

### Supplementary information S6

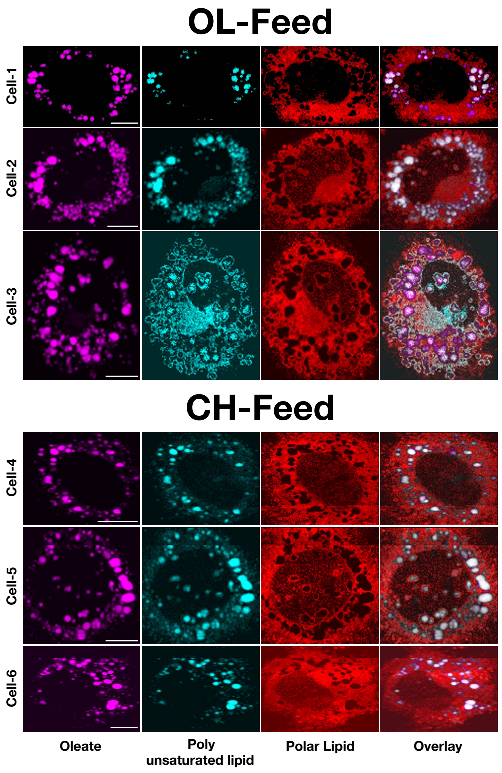


Figure S5(a). Distribution of lipids in HuH7 cells. Scale bar 10 μm.

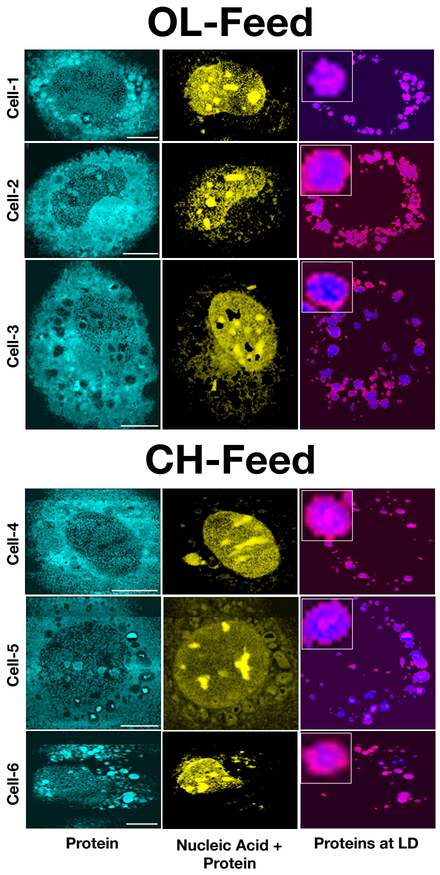


Figure S5(b). Distribution of proteins and nucleic acids in HuH7 cells. Scale bar 10 μm.

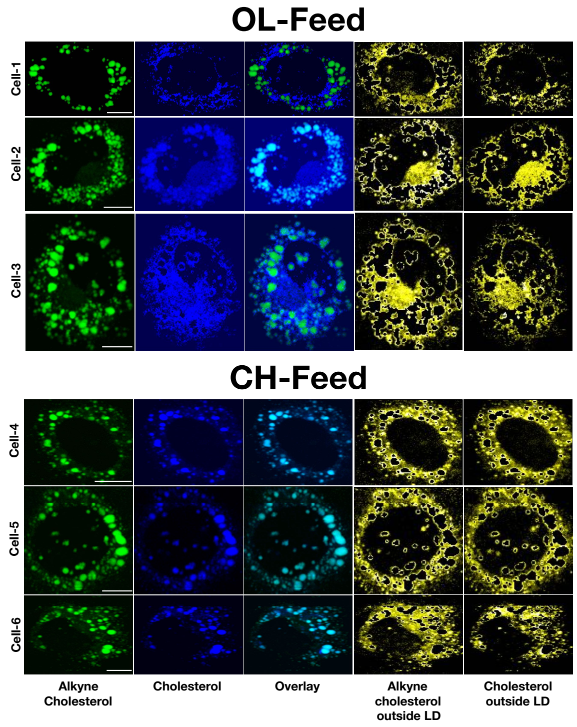


Figure S5(c). Distribution of sterols in HuH7 cells. Scale bar 10 μm.

### Supplementary information S7

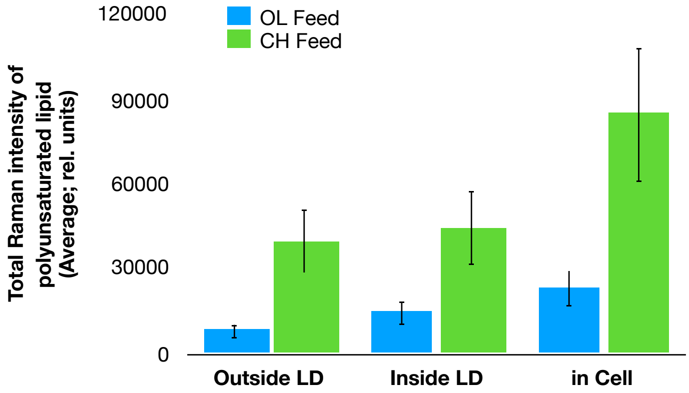


Figure S6. Concentrations of polyunsaturated lipid.
